# The *p* factor: Genetic analyses support a general dimension of psychopathology in childhood and adolescence

**DOI:** 10.1101/591354

**Authors:** A.G. Allegrini, R. Cheesman, K. Rimfeld, S. Selzam, JB. Pingault, T.C. Eley, R. Plomin

## Abstract

**Background:** Diverse behaviour problems in childhood correlate phenotypically, suggesting a general dimension of psychopathology that has been called the p factor. The shared genetic architecture between childhood psychopathology traits also supports a genetic p. This study systematically investigates the manifestation of this common dimension across self-, parent- and teacher-rated measures in childhood and adolescence.

**Methods:** The sample included 7,026 twin pairs from the Twins Early Development Study (TEDS). First, we employed multivariate twin models to estimate common genetic and environmental influences on p based on diverse measures of behaviour problems rated by children, parents and teachers at ages 7, 9, 12 and 16 (depressive symptoms, emotional problems, peer problems, autistic symptoms, hyperactivity, antisocial, conduct and psychopathic symptoms). Second, to assess the stability of genetic and environmental influences on p across time, we conducted longitudinal twin modelling of the first phenotypic principal components of childhood psychopathological measures across each of the four ages. Third, we created a genetic p factor in 7,026 unrelated genotyped individuals based on eight polygenic scores for adult psychiatric disorders to estimate how a general polygenic predisposition to adult psychiatric disorders relates to childhood p.

**Results:** Behaviour problems were consistently correlated phenotypically and genetically across ages and raters. The p factor is substantially heritable (50-60%), and manifests consistently across diverse ages and raters. Genetic correlations of p components across childhood and adolescence suggest stability over time (49-78%). A polygenic general psychopathology factor, derived from studies of adult psychiatric disorders consistently predicted a general phenotypic p factor across development.

**Conclusions:** Diverse forms of psychopathology consistently load on a common p factor, which is highly heritable. There are substantial genetic influences on the stability of p across childhood. Our analyses indicate genetic overlap between general risk for psychiatric disorders in adulthood and p in childhood, even as young as age 7. The p factor has far-reaching implications for genomic research and, eventually, for diagnosis and treatment of behaviour problems.

## Introduction

The p factor, analogous to the concept of general intelligence (‘g’), reflects the observation that individuals who score highly for certain symptoms also score highly on others (Lahey et al. 2012; Caspi et al. 2014). Recent research suggests that this single continuous dimension can, in part, summarise and explain liability to a wide range of psychopathologies in childhood.

Interest in the p factor stemmed initially from high levels of psychopathological comorbidity in adults. The co-occurrence of psychiatric disorders is strikingly high, with up to 50% of individuals diagnosed with a mental illness going on to develop two or more comorbidities in a 12-month period (Kessler et al. 2005). Already during childhood and adolescence, forms of psychopathology are often comorbid. A recent report found that 1 in 20 British young people under 20 years of age met criteria for 2 or more mental disorders (NHS Digital 2017).

Quantitative genetic research suggests that shared genetic factors contribute substantially to the observed co-occurrence of psychopathological traits (Plomin et al. 2016). Several multivariate twin and family studies have replicated the finding that a common genetic factor influences a wide range of emotional and behavioural problems in childhood (Waldman et al. 2016; Tackett et al. 2013; Lahey et al. 2011). Many studies have investigated developmental genetic effects on specific psychopathological traits in childhood (e.g. Pingault et al. 2015), yet little is known about the genetic and environmental architecture of general psychopathology across development. Stability and change in p across time, and the extent to which genetic influences drive age-related patterns remain largely unknown. Here, for the first time, we systematically investigate p across diverse ages, raters and measures in childhood and adolescence.

It is also unknown to what extent a general p factor across earlier development relates to adult psychopathology. In addition to genetic analyses using the twin and family designs, polygenic scores are a new genomic tool that can be used to test for shared genetic effects across traits. Polygenic scores are constructed by aggregating genetic risk across thousands of genetic variants, thus indexing the genetic liability that each individual carries for a specific trait. A landmark study in the field of psychiatric genetics (International Schizophrenia Consortium et al. 2009) first showed that a polygenic score for schizophrenia was also associated with bipolar disorder, suggesting a shared genetic component underlying these two disorders, which has been substantiated further more recently (Cross-Disorder Group of the Psychiatric Genomics Consortium, 2019). Several studies have used polygenic scores for schizophrenia, ADHD, and other psychiatric disorders to predict general psychopathology in childhood. An increasing amount of evidence converges on the finding that few polygenic effects specific to individual aspects of psychopathology remain after conditioning on the p factor (Jones et al. 2018; Jones et al. 2016; Riglin et al. 2018; Brikell et al. 2017). These studies also suggest that genetic risk for psychiatric disorders emerges in childhood, in the form of continuously measured behavior problems. More recently, a study using different genomic methods provided evidence for a ‘polygenic p’ factor, yielding similarly high weights for psychiatric disorders across the different approaches (Selzam et al. 2018). However, no studies to date have empirically related ‘polygenic p’ to ‘phenotypic p’ or systematically tested the architecture of p across development and across different raters.

Here we investigated the structure of general psychopathology across childhood and adolescence. Our study has three aims:

1. Investigate the genetic architecture of p in childhood through common pathway twin models across ages and raters.
2. Test the stability of p across childhood and adolescence through longitudinal quantitative genetic analysis of first principal components of psychopathology across ages (7, 9, 12 and 16) and raters (parent-, teacher- and self-ratings).
3. Estimate associations between childhood phenotypic p and adult polygenic p. The latter can be constructed by principal component analysis of polygenic scores for adult psychiatric disorders created for each TEDS participant.

## Methods

### Sample

The sampling frame is the Twins Early Development Study (TEDS), a multivariate, longitudinal study of >10 000 twin pairs representative of England and Wales, recruited from 1994 - 1996 births (Haworth et al. 2013). Analyses were conducted on a sub-sample of unrelated individuals with available genotype data and their co-twins (N = 7,026). Genomic analyses were limited to unrelated individuals (one twin from each pair).

### Genotyping

Data were available for 3,057 individuals genotyped on the AffymetrixGeneChip 6.0 and 3,969 individuals genotyped HumanOmniExpressExome-8v1.2 arrays. Typical quality control procedures were followed (e.g., samples were removed based on call rate <0.98, MAF <0.5%). Genotypes from the two platforms were separately imputed and then harmonized (for detail see Selzam et al. 2018).

### Measures

Twins Early Development Study (TEDS) measures have been described previously (Haworth et al., 2013). Measures administered at ages 7, 9, 12, and 16 were included in our analyses. Some of these measures (e.g. peer problems, prosocial behaviour (reversed), autistic traits) have not previously been used in other studies of general psychopathology, but we adopted a hypothesis-free approach in an attempt to capture a general trait that is pervasive across diverse domains. For similar reasons, we included all measures available at each age, even though some measures (e.g., aggression) were available only at one age.

For all phenotypes, z-standardised residuals were derived for each scale regressed on sex and age. Composite scores were calculated as unit-weighted means, with the requirement of complete data for at least half the individual measures contributing each composite (i.e., 3 of 4 measures, or 2 of 3, sub-scales measures). All procedures were executed using RStudio (Version 1.1.419; RStudio 2019).

#### Age 7 measures

We used both parent and teacher ratings of all subscales of the Strengths and Difficulties Questionnaire (SDQ) (Hyperactivity, Conduct Problems, Peer Problems, Emotional Problems, and Prosocial (reversed); Goodman 1997), as well as the Antisocial Process Screening Device (APSD) and autistic symptoms (ASD).

#### Age 9 measures

The 5 subscales of the SDQ and ASD were included in the set of self-, parent- and teacher-reported measures. In addition, we used parent- and teacher-rated APSD and aggression (a mean of proactive and reactive scales) measures.

#### Age 12 measures

The 5 subscales of the SDQ, the APSD, and the Childhood Autism Spectrum Test (CAST; Williams et al., 2005) were included in the set of self-, parent- and teacher-reported measures. Parent reports of the Moods and Feelings Questionnaire (MFQ) assessing depressive symptoms, and the Conner’s ADHD behaviours measure were also available.

#### Age 16 measures

The 5 subscales of the SDQ, CAST, MFQ and callous-unemotional measures were available from self reports and teacher reports. Parent-rated data on Conner’s ADHD measure was also included.

### Statistical analyses

#### Common pathway twin models of behaviour problem measures for each rater at each age

To estimate the genetic and environmental influence on phenotypic variance in general psychopathology, and to examine loadings of individual psychopathology measures on p, we conducted multivariate twin model-fitting analyses. In the twin design, differences in within-pair trait correlations for monozygotic (MZ) and dizygotic (DZ) twins are used to estimate genetic, shared environmental, and non-shared environmental effects on traits. Greater MZ than DZ similarity indicates additive genetic influence (A). Within-pair similarity that is not due to genetic factors is attributed to shared environmental influences (C). Non-shared environment (E) accounts for individual-specific factors that influence differences among siblings from the same family, plus measurement error. We considered genetic and environmental associations between all psychopathology measures at each age and separately for each rater. Specifically, we fit the data to the common pathway model (Rijsdijk 2005). This is a multivariate twin model, in which common genetic and environmental variation influence all measures via a single common latent (p) factor. The model allows the estimation of genetic and environmental influences on a common factor (p), and of the factor loadings of each measure of psychopathology on the latent liability (p).

#### Longitudinal twin analysis: Cholesky decomposition of phenotypic principal components

We performed a Cholesky decomposition of the parent-rated phenotypic p principal components, allowing for the investigation of stability and innovation in the genetic and environmental influences on our measures of p across the four ages. We focused on parent-rated data since measures were much more consistent across time than for self report and teacher report. The first genetic factor (A1) represents genetic influences on p at age 7. The extent to which these same genes also influence p at ages 9, 12, and 16 is also estimated, and is represented by the diagonal pathways from A1 to the other variables. The second genetic factor (A2) represents genetic influences on p at age 9 that are independent of those influencing age 7. The extent to which these genes also influence p at ages 12 and 16 is also estimated. The third genetic factor (A3) indexes genetic influences on p at age 12 that are independent of genetic influences shared with the previous ages. The impact of these genes on age 16 general psychopathology is also estimated. Finally, the fourth genetic factor (A4) represents residual genetic influences on age 16 general psychopathology. The same decomposition is done for the shared environmental and non-shared environmental influences (C1–4 and E1–4, respectively). All twin model fitting analysis using full-information maximum likelihood were carried out with structural equation modelling software OpenMx (Neale et al., 2016).

#### Extracting p: Principal Component Analyses (PCA)

In preparation for longitudinal analyses and genomic prediction analyses, we obtained the first principal component (1^st^ PC) of behaviour problem phenotypes at each age separately for child, parent and teacher ratings. Only individuals with complete data were used to generate PCs. We report full results from PCA, which in themselves give insights into the phenotypic architecture of p in childhood. The variance explained by the first PC suggests how much the p factor underpins diverse forms of psychopathology, and loadings of each measure on the first PC indicate the extent to which variables reflect general psychopathology.

We also obtained the first PC from polygenic scores for psychiatric disorders (polygenic p). We used publicly available genome-wide association summary statistics for 8 major psychiatric traits: autism spectrum disorder (Grove et al. 2019), major depressive disorder (MDD; (Wray et al. 2018)), bipolar disorder (BIP); schizophrenia (SCZ; (Pardiñas et al. 2018)); attention deficit hyperactivity disorder (ADHD; (Demontis et al. 2019)); obsessive compulsive disorder (OCD; (International Obsessive Compulsive Disorder Foundation Genetics Collaborative (IOCDF-GC) and OCD Collaborative Genetics Association Studies (OCGAS) 2018)); anorexia nervosa (AN; (Duncan et al. 2017)); post-traumatic stress disorder (PTSD; (Duncan et al. 2017)). For each psychiatric disorder, polygenic scores for each TEDS participant were created in LDpred (Vilhjálmsson et al. 2015), assuming a fraction of causal markers of 1 (analysis steps were analogous to Selzam et al. (2018)).

#### Assessing the association between the polygenic 1^st^ PC and the phenotypic 1st PC across childhood

To assess the extent to which the genetic predisposition for a general psychopathology factor relates to p in childhood, we performed ordinary least square regression analyses of phenotypic p on polygenic p at each age separately by each rater. Age, sex and the first 10 genomic principal components were regressed from all dependent and independent variables, and standardized residuals were used in all linear models.

## Results

### Common pathway twin models

Common pathway twin models showed substantial heritability for the p-factor at each age for all raters (50% to 60%. See Figure 1 for parent-rated measures and Supplementary Figures S1 and S2 for teacher-rated and child-rated measures, respectively. Shared environmental effects were moderate for the parent-rated common factors (~30%) (Supplementary Figure 1), absent for the teacher-rated common factors (~0%; Supplementary Figure S1), and weak for the self-rated common factors (~15%, declining with age; Supplementary Figure S2). Autistic symptoms, conduct problems, antisocial behavior and psychopathic symptoms loaded the highest on the parent-rated and teacher-rated common factors, while emotional problems, depression and anxiety loaded the highest for the child-rated p factor. We also found substantial specific genetic and environmental variance for all measures suggesting unique influences on psychopathological measures beyond the p factor. See Supplementary Table S1 for model-fitting parameters, including sample sizes of measures, which ranged from 2216 to 5592 twin pairs who also had genotype data, and see Supplementary Table S2 for model fit statistics.

**Figure 1:**
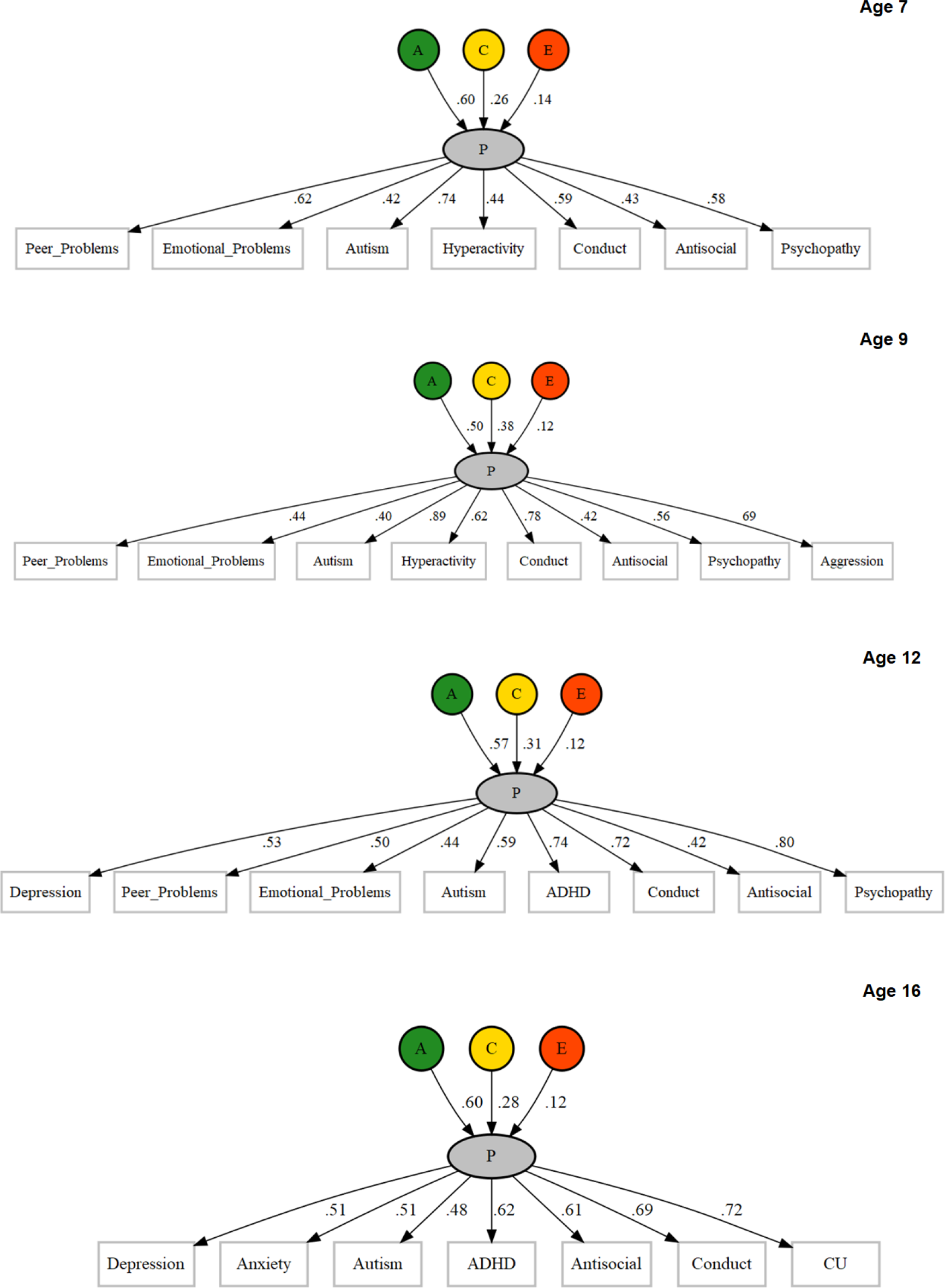
Common pathway twin models of p (parent rated) at ages 7, 9, 12, and 16

### Cholesky decomposition of p across development

The Cholesky decomposition of principal components suggests stability of genetic effects on general psychopathology across childhood and adolescence, in addition to new genetic components at each age, as shown in Figure 2 for parent-ratings. Supplementary Figure S5 shows genetic correlations derived from a correlated factors solution. Age-to-age genetic correlations derived from these results are high, ranging from 0.49 to 0.78 (see Supplementary Figure 5). Supplementary Figures S3 and S4 present the Cholesky model-fitting results for shared and non-shared environmental variance components, respectively. Supplementary Figures S6 to S15 indicates phenotypic correlations among psychopathology measures at all ages and for all raters. Figure S16 shows that correlations between phenotypic principal components across age are also substantial, ranging from .47 to .68. These correlations are notably similar to genetic correlations from the Cholesky model. Supplementary Table S3 lists loadings of observed measures on first principal components, which shows that loadings are consistently substantial for all measures, ages and raters. The first unrotated principal component of phenotypic measures accounted for 40% to 50% of the variance across ages and raters (see Supplementary Table S4, which also shows the sample sizes for each 1^st^ PC, which ranged from 1391 to 4490).

**Figure 2:**
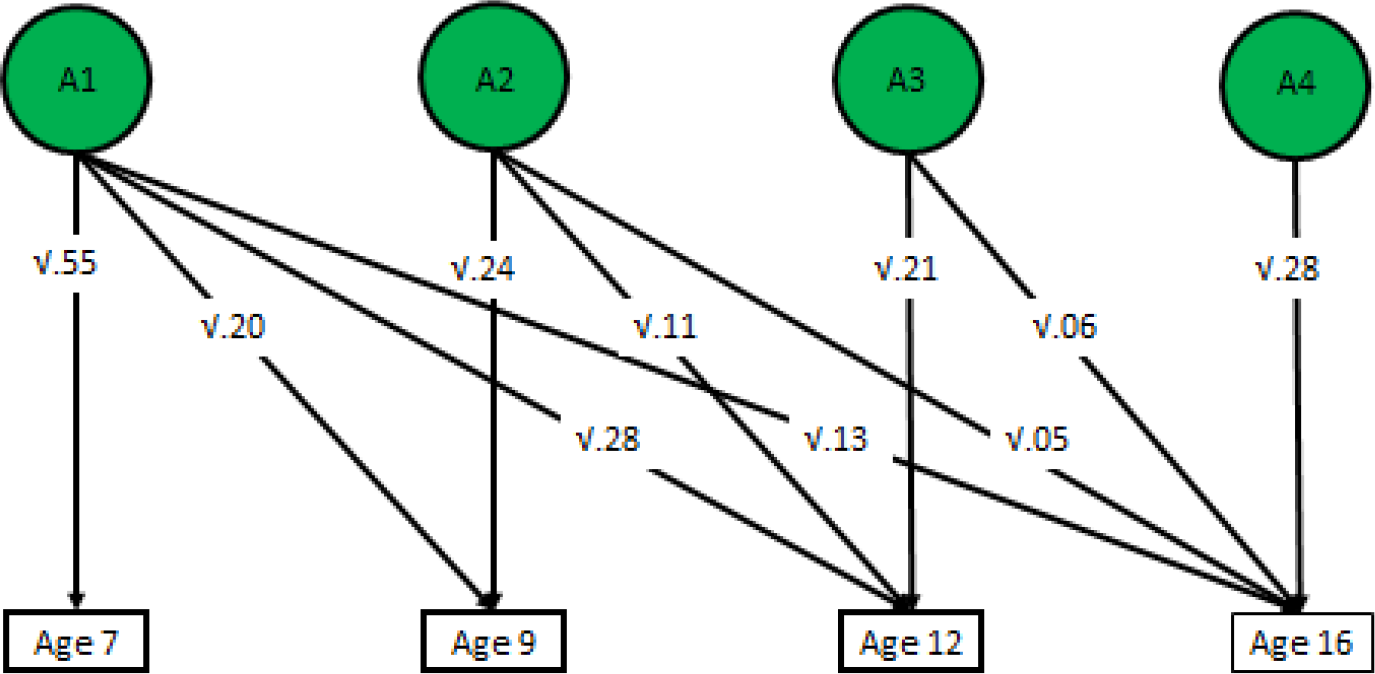
Additive genetic influences on parent-rated p across age, derived from longitudinal twin model-fitting (Cholesky decomposition)

### Prediction of phenotypic p with polygenic p

A polygenic p score defined as the first unrotated principal component of polygenic scores for adult psychiatric disorders was significantly associated with phenotypic p scores in childhood, predicting 0.3% to 0.9% of the variance across ages and raters. See Supplementary Table S5 for full polygenic prediction results. Prediction was generally consistent across ages and raters, although standard errors are largely overlapping (see Figure 3). Supplementary Figure S17 shows correlations between the polygenic scores in TEDS used to derive polygenic p. Although these correlations are modest (0.01 to 0.32), the first principal component of polygenic scores from psychiatric traits explained up to 20% of the polygenic score variability. The pattern of loadings for polygenic p are shown in Supplementary Figure S18.

**Figure 3:**
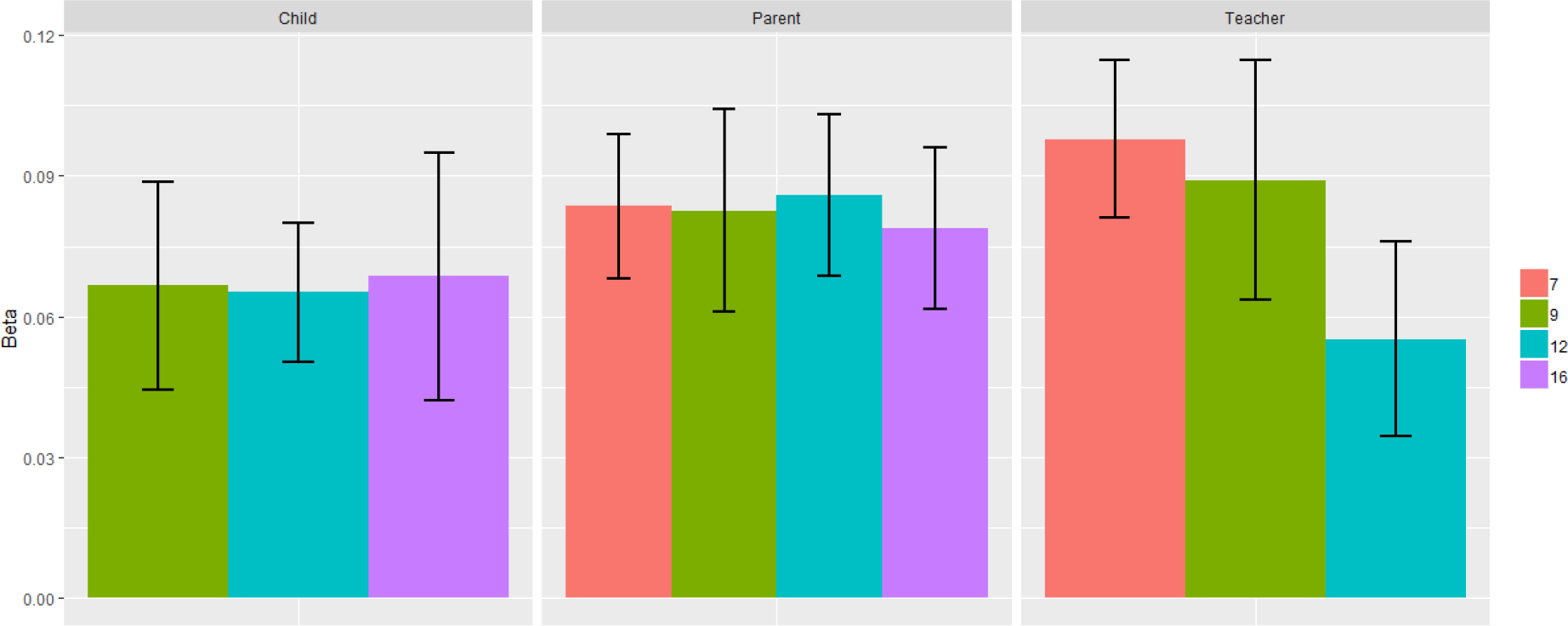
Prediction of phenotypic p with polygenic p by ages and raters

## Discussion

For the first time, we systematically quantified the extent to which a single common factor relates to diverse forms of psychopathology across childhood and adolescence using phenotypic, genetic and genomic methods. Phenotypically, our results confirm previous findings of a strong p factor involving all measures at all ages for all raters. Our genetic results support three main conclusions. Firstly, multivariate twin analyses revealed that 48% to 80% of the variance in the common factor was due to genetic influences, depending on age and raters considered. Secondly, longitudinal twin model-fitting showed that this genetic p factor was stable across time. Thirdly, polygenic prediction analyses demonstrate that there are shared genetic influences connecting childhood psychopathology to general risk for adult psychiatric disorders. In sum, these analyses provide further evidence that a common genetic substrate permeates the landscape of psychopathology, across measures, ages and raters. It is important to note that although we found a consistent and stable genetic p factor across childhood and adolescence, substantial unique genetic and environmental influences indicate that there are also genetic components specific to each trait and each age beyond p.

Our common pathway twin modelling analyses, for which we adopted a hypothesis-free approach to the inclusion of measures, show that diverse psychopathological traits contribute to p. Furthermore, it is commonly acknowledged that all psychopathological traits are dimensional traits both at the phenotypic and genetic levels (Plomin et al. 2009). Future research might investigate the extent to which p might extend to other behavioural domains. For example, suggestive evidence of links between p and personality have begun to emerge (Rosenström et al. 2018).

Differences between raters in our common pathway twin analyses suggested some additional insights. Firstly, inspection of the loadings of psychopathology measures revealed that ‘externalising’ problems relating to conduct and antisocial behavior contributed most to parent- and teacher-rated common factors, whereas ‘internalising’ problems such as depression and anxiety loaded the highest for the child-rated p factor. This could suggest that parents report on overt behaviours, which might stem from worry and sadness from the child’s perspective. Secondly, we observed that shared environmental influences were moderate for the parent-report-based p factor, but negligible for self- and teacher-rated p, respectively. This pattern of results is most likely due to rater bias in that parent ratings are based on a single informant rating both twins, whereas for teacher and self ratings different informants rate each twin (Bartels et al. 2004).

Our longitudinal twin model-fitting and polygenic scoring revealed substantial genetic influences on stability of general psychopathology across childhood. Our polygenic score results suggest that these stable genetic influences overlap with those underlying adult psychiatric disorders. Future research could assess influences on different temporal trajectories of p across childhood and adolescence. One study recently showed that polygenic scores for neurodevelopmental disorders (schizophrenia, ADHD) and depression were associated with early adolescent onset depression, whereas later onset depression was only predicted significantly by depression polygenic scores (Rice et al. 2018). This could be repeated with more powerful polygenic p scores.

Naturally, through the course of multivariate longitudinal studies like TEDS, there are changes in available measures and informants, which in turn can introduce variability in the pattern of results. That is, our measures of p are not perfect indices of general liability to psychopathology, but reflect the specific measures and raters available at each age. This is problematic when estimating genetic and environmental influences on stability and change in p across time. Specifically, any innovation cannot solely be attributed to p, as it will reflect new influences on new measures that were not available at the previous age. This criticism is difficult to overcome even with the availability of consistent data: exactly the same measure at different time points does not necessarily reflect the same thing. We consider that the availability of varied measures is a strength rather than a limitation of the present study because this means that our strong evidence for genetic p and genetic stability for p emerges despite the use of different measures. In the cognitive literature on g, this phenomenon is known as the *indifference of the indicator* – any set of diverse cognitive measures yields a strong g factor (Spearman, 1904). Factor loadings were consistently substantial, not only across measures but also across ages and raters. Importantly, the phenotypic correlations between first principal components across time (ranging between ~0.5 and ~0.7) suggest that p indexes a consistent core psychopathology trait.

The fact that we can predict childhood p using polygenic p derived from adult case-control genome-wide association studies has several interesting implications. Firstly, it suggests that in young children there are already manifestations of genetic risk for adult psychiatric disorders. In other words, early onset behavioural and emotional problems are early signs of psychiatric genetic risk. This is particularly striking given that genetic stability for psychopathology often does not begin until adolescence, and supports other evidence for the usefulness of early intervention for psychiatric problems. The second implication of the genetic overlap between p in childhood and adulthood relates to research design. Specifically, researchers could increase the power of genome-wide association studies to detect DNA variation associated with general risk for psychopathology by aggregating diverse traits across wide age ranges. One way to implement this is a common-factor genome-wide association analysis using Genomic SEM (Grotzinger et al. 2018). Similarly, the modest power of psychiatric polygenic scores to predict traits in childhood could be enhanced using multi-trait frameworks to generate predictors that leverage the shared genetic risk between traits (e.g. SMTpred; Maier et al. 2018).

The current clinical zeitgeist focuses on specificity. The recognition that a common factor transcends diverse aspects of psychopathology in childhood is of primary importance, as this knowledge can inform early detection of children at risk in the general population.

## Key points

*We investigated the underlying structure of p across diverse measures, ages and raters, and consistently found a substantial genetic component, in line with previous theory.

*We showed that this genetic component is stable across time, with influences in childhood being pervasive across development through to adolescence.

*Genomic analyses revealed shared genetic risk between p in children as young as 7 and general risk for adult psychiatric disorders.

*We provide further evidence that, in addition to residual variation specific to each trait, a common genetic substrate permeates the landscape of psychopathology.

## Acknowledgements

We gratefully acknowledge the ongoing contribution of the participants in the Twins Early Development Study (TEDS) and their families. TEDS is supported by a program grant to R.P. from the UK Medical Research Council (MR/M021475/1 and previously G0901245), with additional support from the US National Institutes of Health (AG046938). The research leading to these results has also received funding from the European Research Council under the European Union’s Seventh Framework Programme (FP7/2007–2013)/ grant agreement no. 602768 and ERC grant agreement no. 295366). R.P. is supported by a Medical Research Council Professorship award (G19/2). T.C. Eley is part funded by the above program grant from the UK Medical Research Council (MR/M021475/1). This study represents independent research part funded by the National Institute for Health Research (NIHR) Biomedical Research Centre at South London and Maudsley NHS Foundation Trust and King’s College London. The views expressed are those of the author(s) and not necessarily those of the NHS, the NIHR or the Department of Health. High-performance computing facilities were funded with capital equipment grants from the GSTT Charity (TR130505) and Maudsley Charity (980). R.C. is supported by an ESRC studentship. AGA has received funding from the European Union’s Horizon 2020 research and innovation programme under the Marie Sklodowska-Curie grant agreement no. 721567. J.-B.P. is a fellow of MQ: Transforming Mental Health (MQ16IP16).

## Abbreviations

P: general psychopathology factor
PCA: principal component analysis

## References

Bartels, M., Boomsma, D.I., Hudziak, J.J., Rietveld, M.J., van Beijsterveldt, T.C. and van den Oord, E.J., 2004. Disentangling genetic, environmental, and rater effects on internalizing and externalizing problem behavior in 10-year-old twins. Twin Research and Human Genetics, 7(2), pp.162–175.

Boker, S., Neale, M., Maes, H., Wilde, M., Spiegel, M., Brick, T., Spies, J., Estabrook, R., Kenny, S., Bates, T., Mehta, P. and Fox, J. 2011. Openmx: an open source extended structural equation modeling framework. Psychometrika 76(2), pp.306–317.

Brikell, I., Larsson, H., Lu, Y., Pettersson, E., Chen, Q., Kuja-Halkola, R., Karlsson, R., Lahey, B.B., Lichtenstein, P. and Martin, J. 2017. The contribution of common genetic risk variants for ADHD to a general factor of childhood psychopathology. BioRxiv.

Caspi, A., Houts, R.M., Belsky, D.W., Goldman-Mellor, S.J., Harrington, H., Israel, S., Meier, M.H., Ramrakha, S., Shalev, I., Poulton, R. and Moffitt, T.E. 2014. The p Factor: One General Psychopathology Factor in the Structure of Psychiatric Disorders? Clinical psychological science: a journal of the Association for Psychological Science 2(2), pp. 119–137.

Cheesman, R., Selzam, S., Ronald, A., Dale, P.S., McAdams, T.A., Eley, T.C. and Plomin, R. 2017. Childhood behaviour problems show the greatest gap between DNA-based and twin heritability.Translational Psychiatry.

Cross-Disorder Group of the Psychiatric Genomics Consortium, Lee, P. H., Anttila, V., Won, H., Feng, Y.-C. A., Rosenthal, J., et al. (2019). Genome wide meta-analysis identifies genomic relationships, novel loci, and pleiotropic mechanisms across eight psychiatric disorders. bioRxiv, 528117. http://doi.org/10.1101/528117

Demontis, D., Walters, R.K., Martin, J., Mattheisen, M., Als, T.D., Agerbo, E., Baldursson, G., Belliveau, R., Bybjerg-Grauholm, J., Bækvad-Hansen, M., Cerrato, F., Chambert, K., Churchhouse, C., Dumont, A., Eriksson, N., Gandal, M., Goldstein, J.I., Grasby, K.L., Grove, J., Gudmundsson, O.O. and Neale, B.M. 2019. Discovery of the first genome-wide significant risk loci for attention deficit/hyperactivity disorder. Nature Genetics 51(1), pp. 63–75.

Duncan, L., Yilmaz, Z., Gaspar, H., Walters, R., Goldstein, J., Anttila, V., Bulik-Sullivan, B., Ripke, S., Eating Disorders Working Group of the Psychiatric Genomics Consortium, Thornton, L., Hinney, A., Daly, M., Sullivan, P.F., Zeggini, E., Breen, G. and Bulik, C.M. 2017. Significant Locus and Metabolic Genetic Correlations Revealed in Genome-Wide Association Study of Anorexia Nervosa.The American Journal of Psychiatry 174(9), pp. 850–858.

Goodman, R. 1997. The Strengths and Difficulties Questionnaire: A Research Note. Journal of Child Psychology and Psychiatry 38(5), pp. 581–586.

Grotzinger, A.D., Rhemtulla, M., de Vlaming, R., Ritchie, S.J., Mallard, T.T., Hill, W.D., Ip, H.F., McIntosh, A.M., Deary, I.J., Koellinger, P.D., Harden, K.P., Nivard, M.G. and Tucker-Drob, E.M. 2018. Genomic SEM Provides Insights into the Multivariate Genetic Architecture of Complex Traits. BioRxiv.

Grove, J., Ripke, S., Als, T.D., Mattheisen, M., Walters, R.K., Won, H., Pallesen, J., Agerbo, E., Andreassen, O.A., Anney, R., Awashti, S., Belliveau, R., Bettella, F., Buxbaum, J.D., Bybjerg-Grauholm, J., Bækvad-Hansen, M., Cerrato, F., Chambert, K., Christensen, J.H., Churchhouse, C. and Børglum,A.D. 2019. Identification of common genetic risk variants for autism spectrum disorder.Nature Genetics 51(3), pp. 431–444.

Haworth, C.M.A., Davis, O.S.P. and Plomin, R. 2013. Twins Early Development Study (TEDS): a genetically sensitive investigation of cognitive and behavioral development from childhood to young adulthood. Twin Research and Human Genetics 16(1), pp. 117–125.

International Obsessive Compulsive Disorder Foundation Genetics Collaborative (IOCDF-GC) and OCD Collaborative Genetics Association Studies (OCGAS) 2018. Revealing the complex genetic architecture of obsessive-compulsive disorder using meta-analysis. Molecular Psychiatry 23(5), pp. 1181–1188.

International Schizophrenia Consortium, Purcell, S.M., Wray, N.R., Stone, J.L., Visscher, P.M.,O’Donovan, M.C., Sullivan, P.F. and Sklar, P. 2009. Common polygenic variation contributes to risk of schizophrenia and bipolar disorder. Nature 460(7256), pp. 748–752.

Jones, H.J., Heron, J., Hammerton, G., Stochl, J., Jones, P.B., Cannon, M., Smith, G.D., Holmans, P., Lewis, G., Linden, D.E.J., O’Donovan, M.C., Owen, M.J., Walters, J., Zammit, S. and 23 and Me Research Team 2018. Investigating the genetic architecture of general and specific psychopathology in adolescence. Translational psychiatry 8(1), p. 145.

Jones, H.J., Stergiakouli, E., Tansey, K.E., Hubbard, L., Heron, J., Cannon, M., Holmans, P., Lewis, G., Linden, D.E.J., Jones, P.B., Davey Smith, G., O’Donovan, M.C., Owen, M.J., Walters, J.T. and Zammit, S. 2016. Phenotypic manifestation of genetic risk for schizophrenia during adolescence in the general population. JAMA psychiatry 73(3), pp. 221–228.

Kessler, R.C., Berglund, P., Demler, O., Jin, R., Merikangas, K.R. and Walters, E.E. 2005. Lifetime prevalence and age-of-onset distributions of DSM-IV disorders in the National Comorbidity Survey Replication. Archives of General Psychiatry 62(6), pp. 593–602.

Lahey, B. B., Applegate, B., Hakes, J. K., Zald, D. H., Hariri, A. R., & Rathouz, P. J. (2012). Is there a general factor of prevalent psychopathology during adulthood?. Journal of abnormal psychology, 121(4), 971.

Lahey, B.B., Van Hulle, C.A., Singh, A.L., Waldman, I.D. and Rathouz, P.J. 2011. Higher-order genetic and environmental structure of prevalent forms of child and adolescent psychopathology. Archives of General Psychiatry 68(2), pp. 181–189.

Maier, R.M., Zhu, Z., Lee, S.H., Trzaskowski, M., Ruderfer, D.M., Stahl, E.A., Ripke, S., Wray, N.R., Yang, J., Visscher, P.M. and Robinson, M.R. 2018. Improving genetic prediction by leveraging genetic correlations among human diseases and traits. Nature Communications 9(1), p. 989.

Neale, M.C., Hunter, M.D., Pritikin, J.N., Zahery, M., Brick, T.R., Kirkpatrick, R.M., Estabrook, R., Bates, T.C., Maes, H.H. and Boker, S.M., 2016. OpenMx 2.0: Extended structural equation and statistical modeling. Psychometrika, 81(2), pp.535–549.

NHS Digital 2017. Mental Health of Children and Young People in England, 2017 [PAS] – NHS Digital [Online]. Available at: https://digital.nhs.uk/data-and-information/publications/statistical/mental-health-of-children-and-young-people-in-england/2017/2017 [Accessed: 11 December 2018].

Pardiñas, A.F., Holmans, P., Pocklington, A.J., Escott-Price, V., Ripke, S., Carrera, N., Legge, S.E., Bishop, S., Cameron, D., Hamshere, M.L., Han, J., Hubbard, L., Lynham, A., Mantripragada, K., Rees, E., MacCabe, J.H., McCarroll, S.A., Baune, B.T., Breen, G., Byrne, E.M. and CRESTAR Consortium 2018. Common schizophrenia alleles are enriched in mutation-intolerant genes and in regions under strong background selection. Nature Genetics 50(3), pp. 381–389.

Pettersson, E., Larsson, H. and Lichtenstein, P. 2016. Common psychiatric disorders share the same genetic origin: a multivariate sibling study of the Swedish population. Molecular Psychiatry 21(5), pp. 717–721.

Pingault, J.-B., Viding, E., Galéra, C., Greven, C.U., Zheng, Y., Plomin, R. and Rijsdijk, F. 2015. Genetic and Environmental Influences on the Developmental Course of Attention-Deficit/Hyperactivity Disorder Symptoms From Childhood to Adolescence. JAMA psychiatry 72(7), pp. 651–658.

Plomin, R., DeFries, J.C., Knopik, V.S. and Neiderhiser, J.M. 2016. Top 10 replicated findings from behavioral genetics. Perspectives on psychological science : a journal of the Association for Psychological Science 11(1), pp. 3–23.

Plomin, R., Haworth, C.M.A. and Davis, O.S.P. 2009. Common disorders are quantitative traits. Nature Reviews. Genetics 10(12), pp. 872–878.

Rice, F., Riglin, L., Thapar, A.K., Heron, J., Anney, R., O’Donovan, M.C. and Thapar, A. 2018. Characterizing Developmental Trajectories and the Role of Neuropsychiatric Genetic Risk Variants in Early-Onset Depression. JAMA psychiatry.

Riglin, L., Thapar, A.K., Leppert, B., Martin, J., Richards, A., Anney, R., Davey Smith, G., Tilling, K., Stergiakouli, E., Lahey, B.B., O’Donovan, M.C., Collishaw, S. and Thapar, A. 2018. The contribution of psychiatric risk alleles to a general liability to psychopathology in early life. BioRxiv.

Rijsdijk, F. 2005. Common Pathway Model. In: Everitt, B. S. and Howell, D.C. eds. Encyclopedia of statistics in behavioral science. Chichester, UK: John Wiley & Sons, Ltd.

Rosenström, T., Gjerde, L.C., Krueger, R.F., Aggen, S.H., Czajkowski, N.O., Gillespie, N.A., Kendler, K.S., Reichborn-Kjennerud, T., Torvik, F.A. and Ystrom, E. 2018. Joint factorial structure of psychopathology and personality. Psychological Medicine, pp. 1–10.

Rstudio 2019. Open source and enterprise-ready professional software for data science - RStudio [Online]. Available at: https://www.rstudio.com/ [Accessed: 15 March 2019].

Selzam, S., Coleman, J.R.I., Caspi, A., Moffitt, T.E. and Plomin, R. 2018. A polygenic p factor for major psychiatric disorders. Translational psychiatry 8(1), p. 205.

Spearman, C.E. (1904). “‘General intelligence’, Objectively Determined And Measured” (PDF). American Journal of Psychology. 15 (2): 201–293. doi:10.2307/1412107.JSTOR1412107

Tackett, J.L., Lahey, B.B., van Hulle, C., Waldman, I., Krueger, R.F. and Rathouz, P.J. 2013. Common genetic influences on negative emotionality and a general psychopathology factor in childhood and adolescence. Journal of Abnormal Psychology 122(4), pp. 1142–1153.

Vilhjálmsson, B.J., Yang, J., Finucane, H.K., Gusev, A., Lindström, S., Ripke, S., Genovese, G., Loh, P.-R., Bhatia, G., Do, R., Hayeck, T., Won, H.-H., Schizophrenia Working Group of the Psychiatric Genomics Consortium, Discovery, Biology, and Risk of Inherited Variants in Breast Cancer (DRIVE) study, Kathiresan, S., Pato, M., Pato, C., Tamimi, R., Stahl, E., Zaitlen, N., Pasaniuc, B. and Price, A.L. 2015. Modeling linkage disequilibrium increases accuracy of polygenic risk scores. American Journal of Human Genetics 97(4), pp. 576–592.

Waldman, I.D., Poore, H.E., van Hulle, C., Rathouz, P.J. and Lahey, B.B. 2016. External validity of a hierarchical dimensional model of child and adolescent psychopathology: Tests using confirmatory factor analyses and multivariate behavior genetic analyses. Journal of Abnormal Psychology 125(8), pp. 1053–1066.

Williams, J., Scott, F., Stott, C., Allison, C., Bolton, P., Baron-Cohen, S. and Brayne, C., 2005. The CAST (childhood asperger syndrome test) test accuracy. Autism, 9(1), pp.45–68.

Wray, N.R., Ripke, S., Mattheisen, M., Trzaskowski, M., Byrne, E.M., Abdellaoui, A., Adams, M.J., Agerbo, E., Air, T.M., Andlauer, T.M.F., Bacanu, S.-A., Bækvad-Hansen, M., Beekman, A.F.T., Bigdeli, T.B., Binder, E.B., Blackwood, D.R.H., Bryois, J., Buttenschøn, H.N., Bybjerg-Grauholm, J., Cai, N. and et al. 2018. Genome-wide association analyses identify 44 risk variants and refine the genetic architecture of major depression. Nature Genetics 50(5), pp. 668–681.

