## Supplementary Information for "The *p* factor: Genetic analyses support a general dimension of psychopathology in childhood and adolescence"

Contents:

Supplementary Figures

- Supplementary Figures S1-S2: Common pathway twin models for child-rated and teacher-rated psychopathology measures by age.
- Supplementary Figures S3-S4: Shared environmental and non-shared environmental influences on p (parent-rated) across age, derived from longitudinal twin model-fitting (Cholesky decomposition)
- Supplementary Figure S5: Correlated factor solution of the longitudinal Cholesky decomposition
- Supplementary Figures S6 to S15: Phenotypic correlations among psychopathology measures used to construct phenotypic p factors.
- Supplementary Figure S16: Correlations of 1st PCs across ages.
- Supplementary Figure S17: Correlations between polygenic scores for psychiatric traits used to construct polygenic p.
- Supplementary Figure S18: PCA results for polygenic p-factor.

Supplementary Tables

- Supplementary Table S1: Additional parameters derived from common pathway twin models of childhood psychopathology in TEDS
- Supplementary Table S2: Model fit statistics for common pathway twin models of childhood psychopathology in TEDS.
- Supplementary Table S3: Loadings on first principal components of psychopathology measures for each age and rater.
- Supplementary Table S4*:* Variance explained by 1st PCs for each age and rater.
- Supplementary Table S5. Association statistics for polygenic p across phenotypic p measures.

Supplementary Figures S1-S2: Common pathway twin models for teacher-rated and child-rated measures by age. *Note: The ACE variance decomposition results for the common factor are presented in the top half of each figure, and the factor loadings of observed psychopathology variables on p are presented in the bottom half. See Supplementary Tables 1 and 2 for additional model parameters and model fit statistics, respectively. Also note that Antisocial = prosocial SDQ scale reversed; and psychopathy = APSD scale.*

*Figure S1. Teacher-rated p factor.*

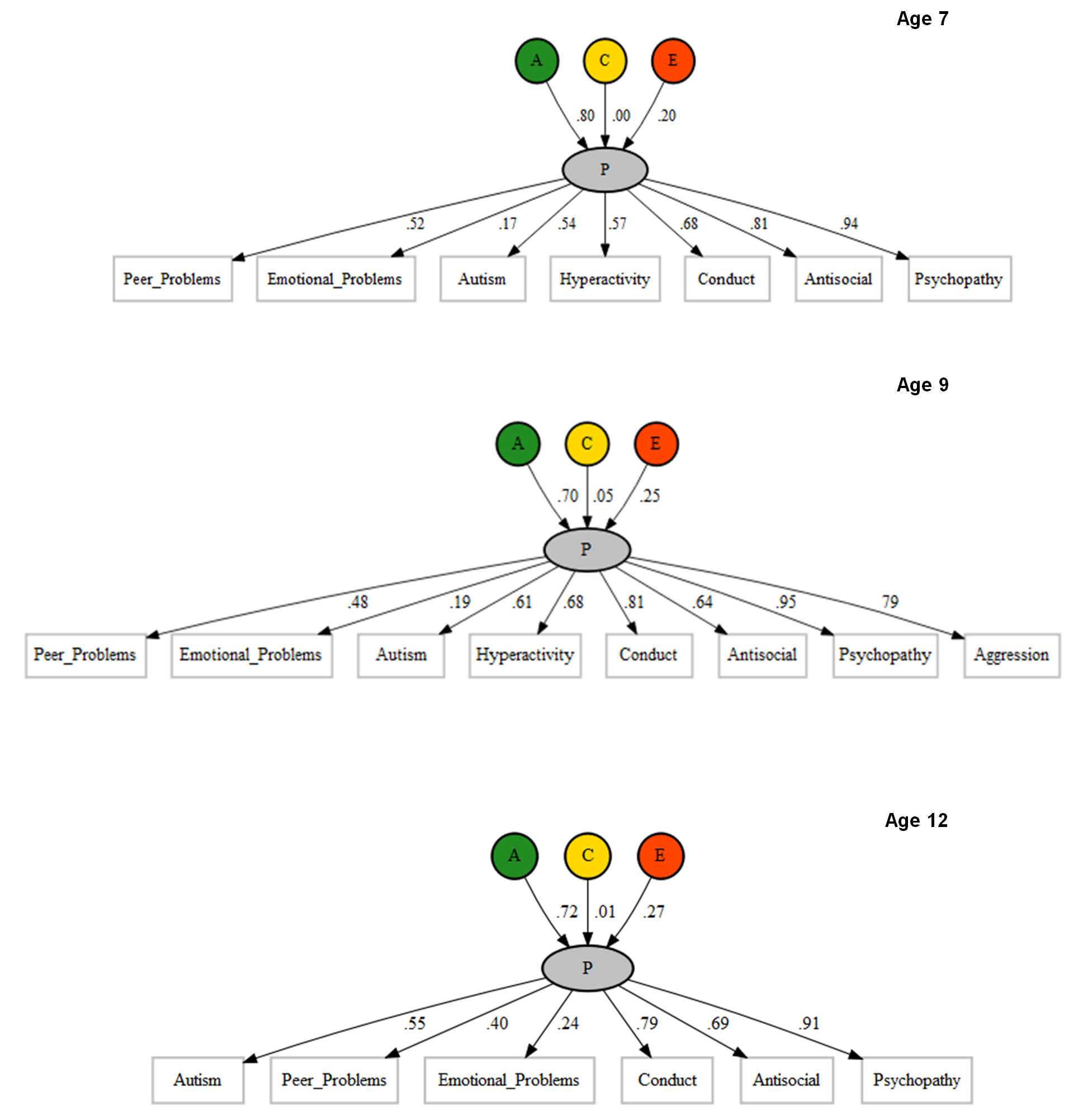

*Figure S2. Child-rated p factor.*

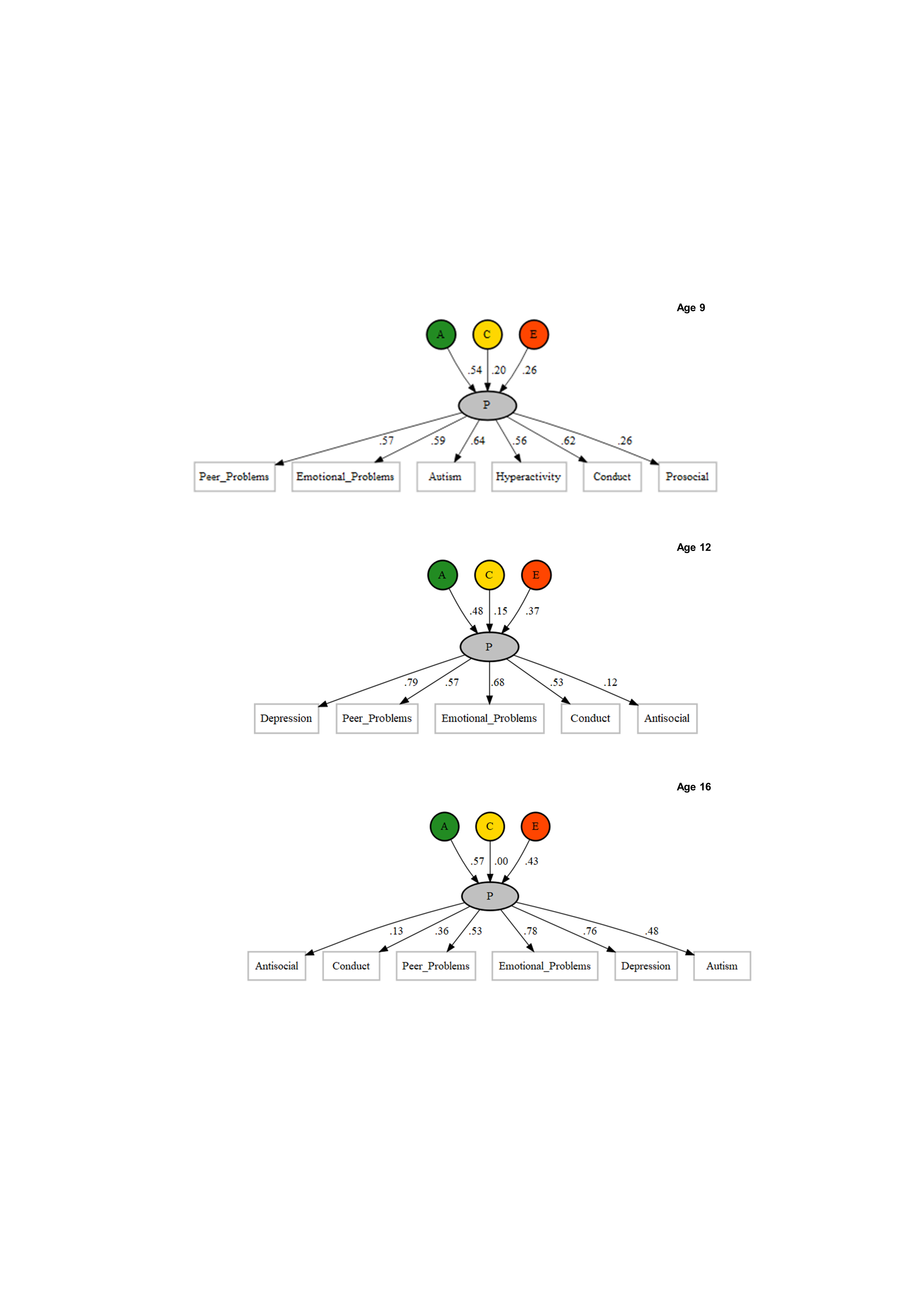

*Figure S3. Shared environmental influences on p (parent-rated) across age, derived from longitudinal twin model-fitting (Cholesky decomposition)*

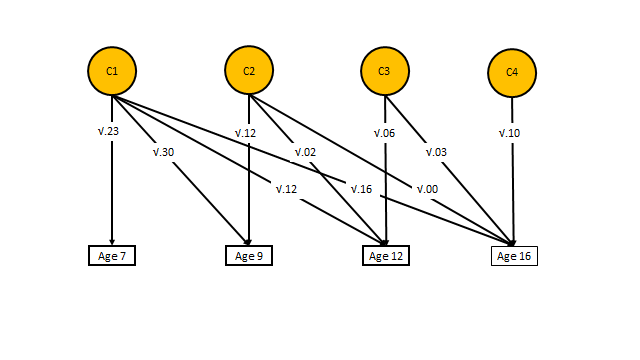

*Figure S4. Unique environmental influences on p (parent-rated) across age, derived from longitudinal twin model-fitting (Cholesky decomposition)*

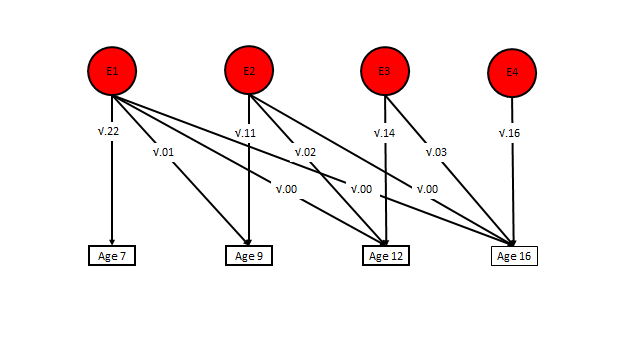

Figure S5. *Correlated factor solution of the longitudinal Cholesky decomposition. The figure shows genetic correlations between and univariate heritability of phenotypic p, defined by the first unrotated principal component of psychopathology measures, using parent-reported data at age 7, 9, 12 and 16.*
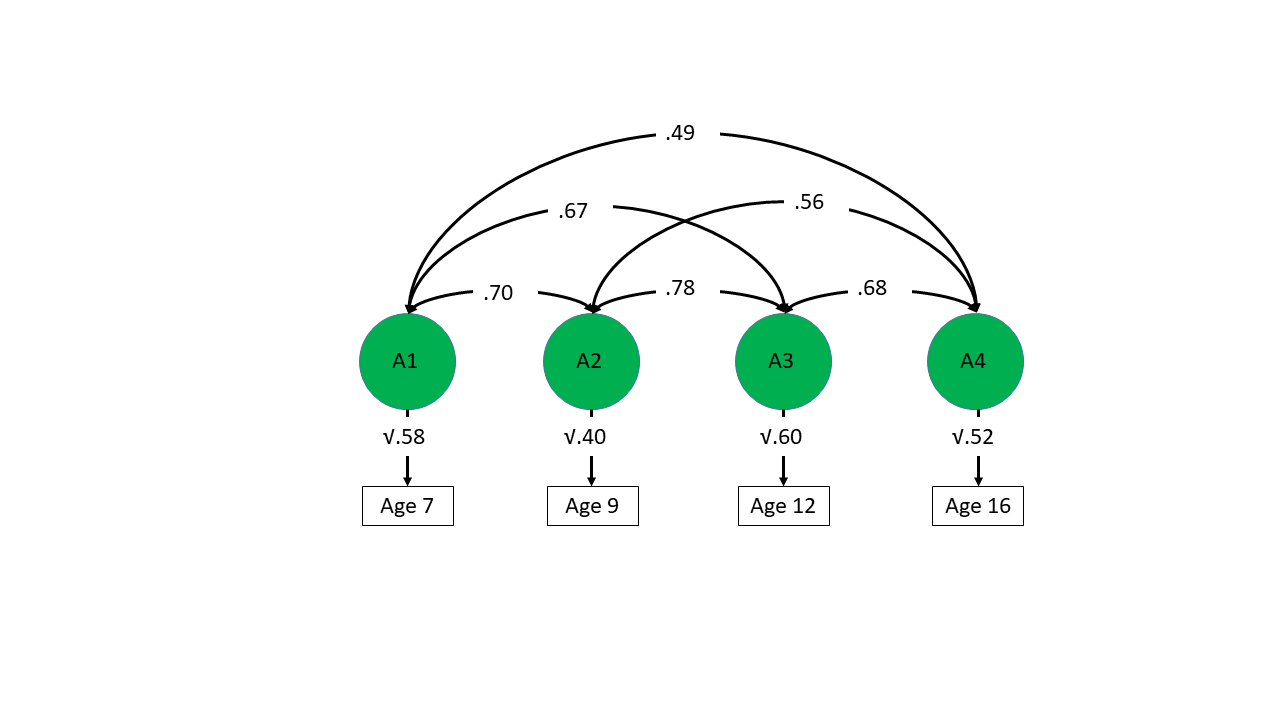

Supplementary Figures S6- to S15: Phenotypic correlations among psychopathology measures used to construct phenotypic p factors. *Darker blue indicates stronger positive correlation.*

*Figure S6. Phenotypic correlations between parent-rated traits at age 7.*

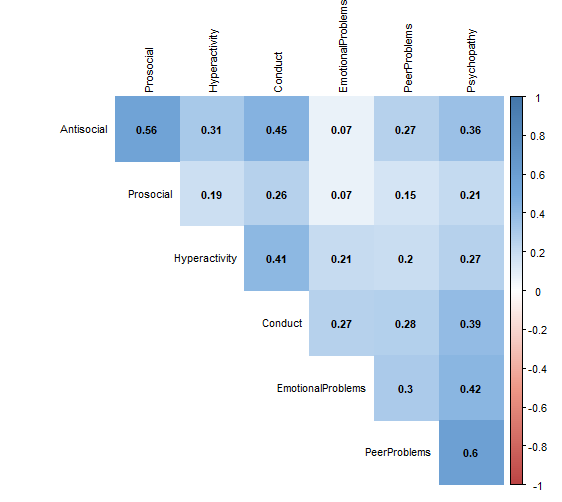

*Figure S7. Phenotypic correlations between Teacher rated psychiatric traits at age 7.*

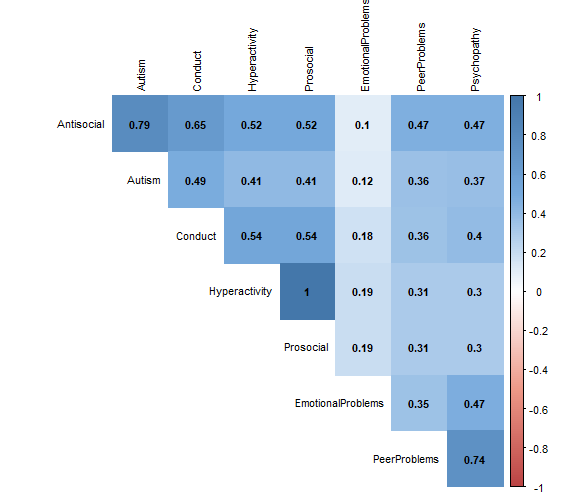

*Figure S8. Phenotypic correlations between Child rated psychiatric traits at age 9.*

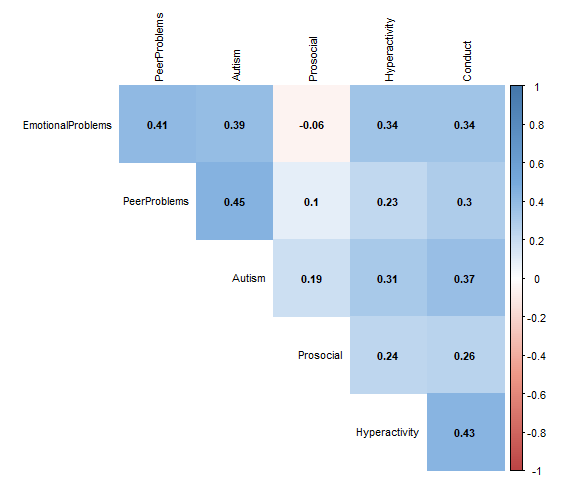

*Figure S9. Phenotypic correlations between Parent rated psychiatric traits at age 9.*

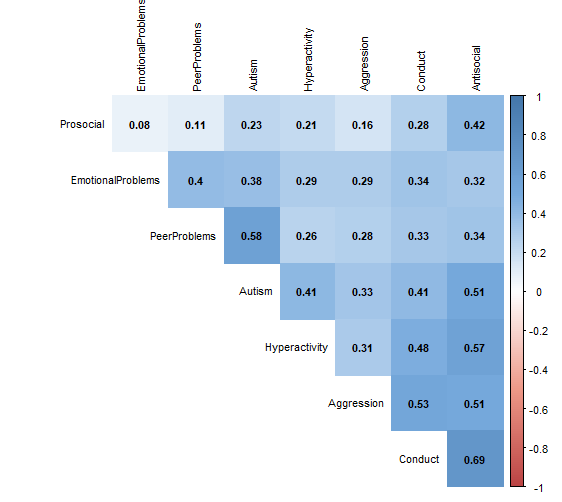

*Figure S10. Phenotypic correlations between Teacher rated psychiatric traits at age 9.*

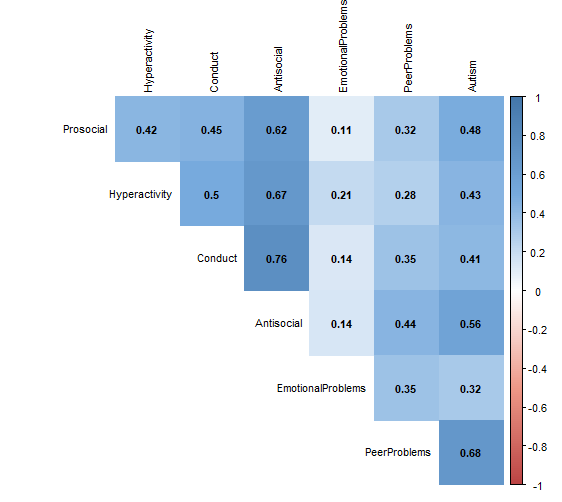

*Figure S11. Phenotypic correlations between Parent rated psychiatric traits at age 12.*

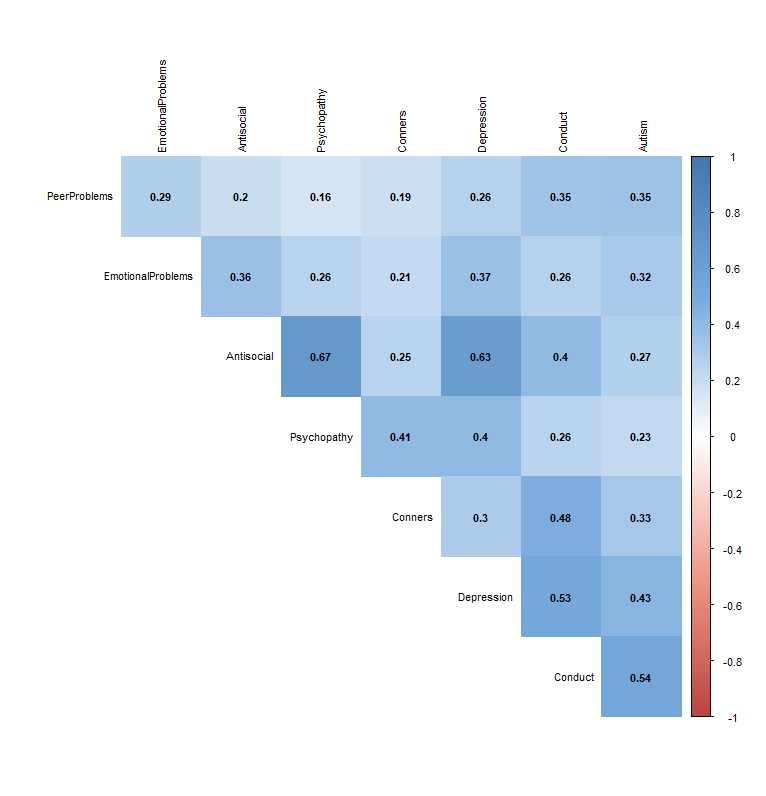

*Figure S12. Phenotypic correlations between Teacher rated psychiatric traits at age 12.*

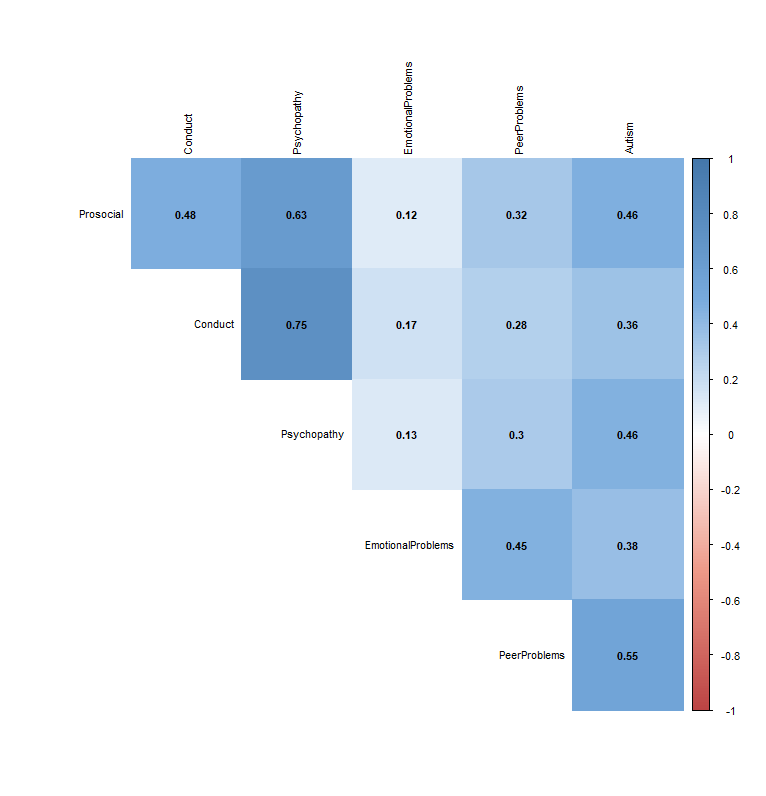

*Figure S13. Phenotypic correlations between Child rated psychiatric traits at age 12.*

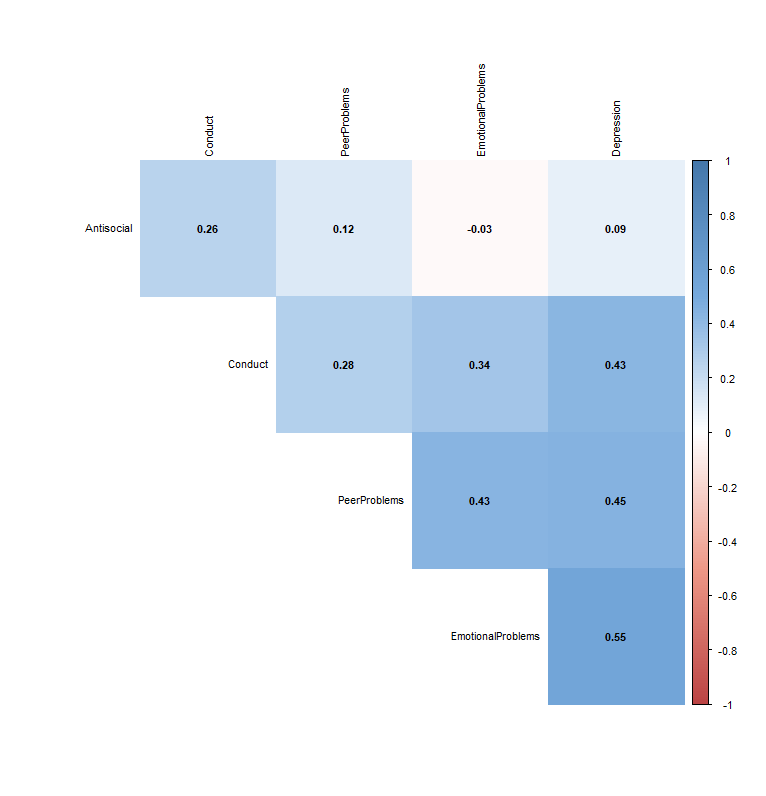

*Figure S14. Phenotypic correlations between Parent rated psychiatric traits at age 16.*

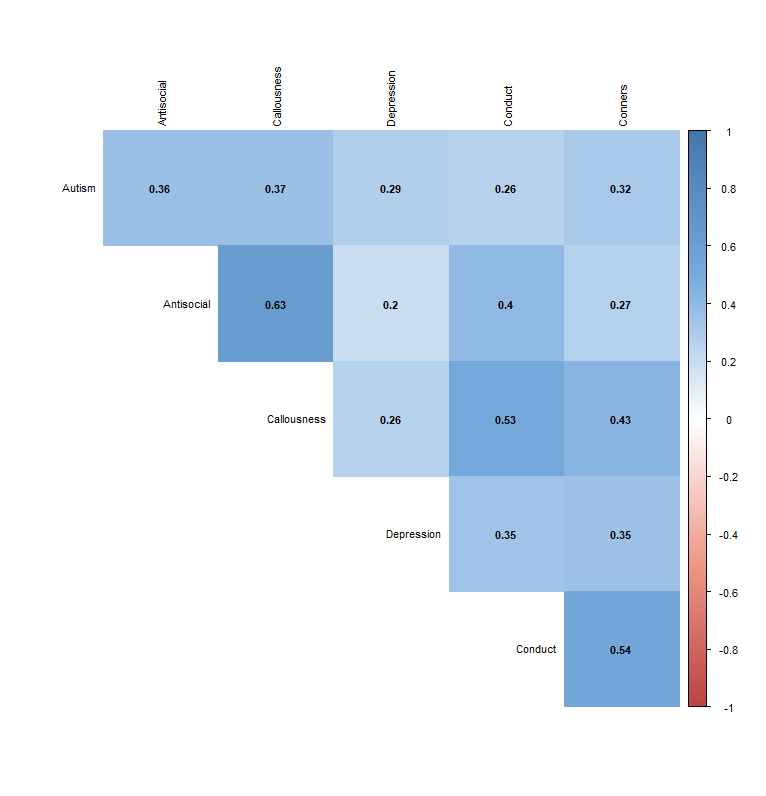

*Figure S15. Phenotypic correlations between Child rated psychiatric traits at age 16.*
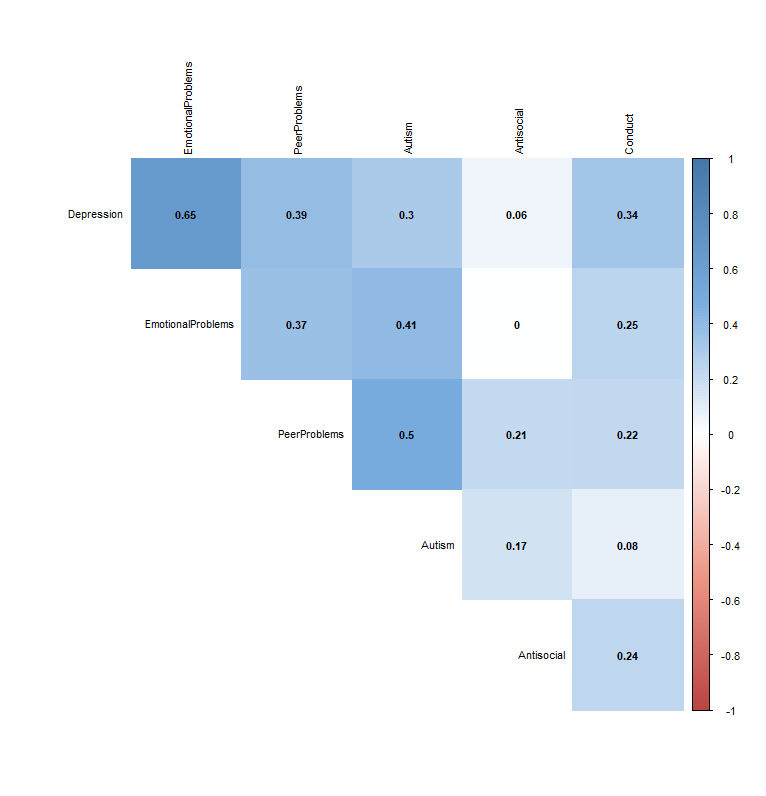

Figure S16. Correlations of 1^st^ PCs of parent-rated measures across ages. Note variable names denote first principal component for parent reported data at age 7 to 16. E.g. P7pc1 = First principal component for parent rated age 7 data.

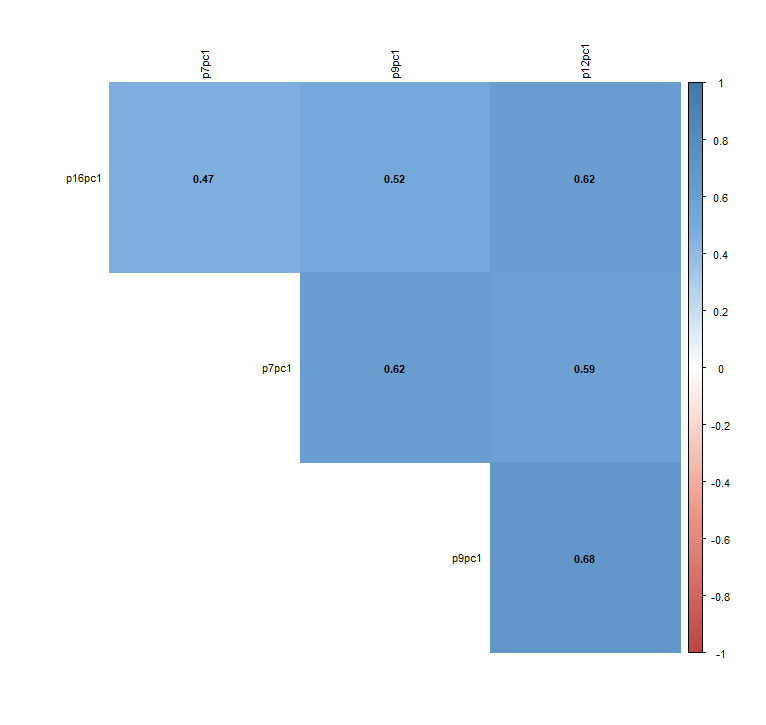

Supplementary Figure S17. Correlations between polygenic scores for psychiatric traits used to construct polygenic p. *OCD =obsessive compulsive disorder; BIP =bipolar disorder; SCZ = schizophrenia; PTSD = Post-traumatic stress disorder; AN = anorexia nervosa; MDD = major depressive disorder; ADHD = attention deficit hyperactivity disorder; AUT = Autism. Darker blue indicates stronger positive correlation.*

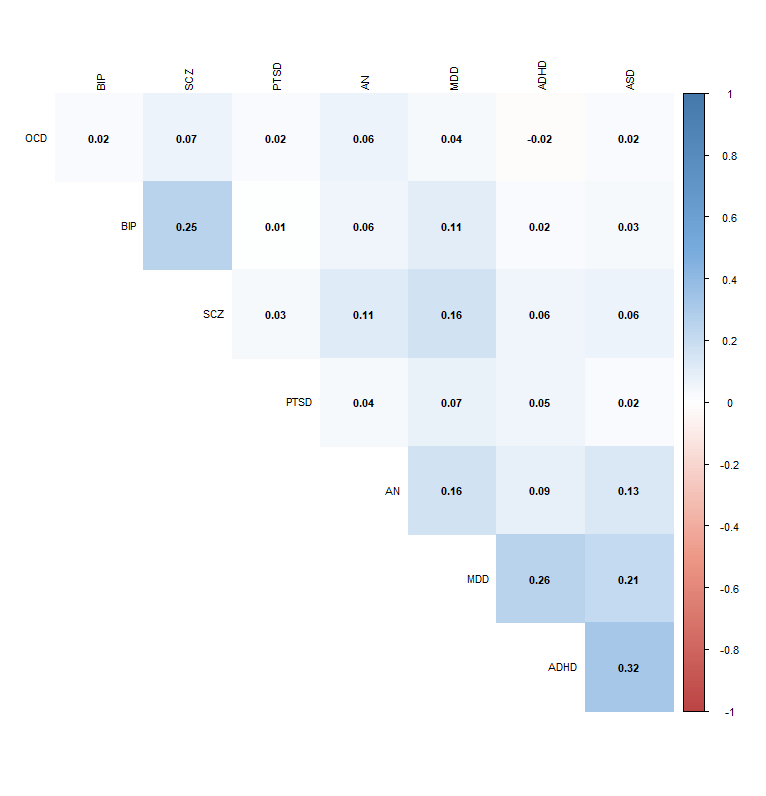

*Supplementary Figure S18. PCA results for polygenic p factor. Note labels for polygenic scores: MDD= major depressive disorder, BIP= bipolar disorder, SCZ= schizophrenia, ASD= autism, AN= anorexia nervosa, ADHD=attention-deficit hyperactivity disorder, OCD= obsessive compulsive disorder, PTSD= post-traumatic stress disorder.*

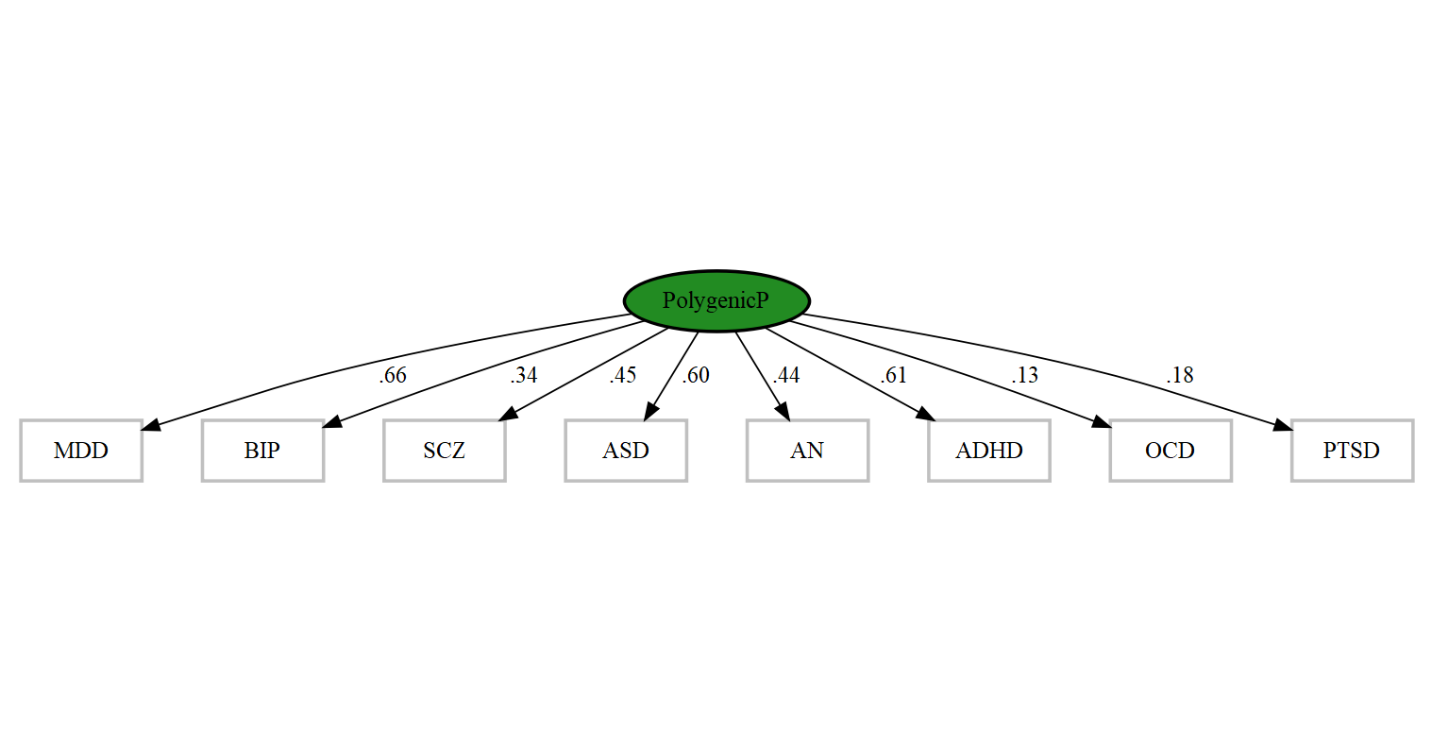

Supplementary Table S1: Additional parameters derived from common pathway twin models of childhood psychopathology. Note that c= common; s=specific; Tot=total.

| **Parent Report Age 7** |  |  |  |  |  |  |  |  |  |  |
| --- | --- | --- | --- | --- | --- | --- | --- | --- | --- | --- |
|  | Ac | As | Cc | Cs | Ec | Es | Tot-H2 | Tot-C2 | Tot-E2 | N |
| Hyperactivity SDQ | 0.11 | 0.23 | 0.05 | 0 | 0.03 | 0.58 | 0.35 | 0.05 | 0.61 | 5588 |
| Conduct SDQ | 0.21 | 0.45 | 0.09 | 0 | 0.05 | 0.2 | 0.66 | 0.09 | 0.25 | 5592 |
| Peer Problems SDQ | 0.23 | 0.29 | 0.1 | 0 | 0.06 | 0.32 | 0.53 | 0.1 | 0.37 | 5591 |
| Emotional Problems SDQ | 0.1 | 0.44 | 0.04 | 0 | 0.02 | 0.39 | 0.54 | 0.04 | 0.41 | 5591 |
| APSD Psychopathy= Narcissism + CU-Traits | 0.2 | 0.36 | 0.09 | 0 | 0.05 | 0.3 | 0.56 | 0.09 | 0.35 | 5587 |
| Asd = Social + Nonsocial | 0.33 | 0.29 | 0.14 | 0 | 0.08 | 0.16 | 0.62 | 0.14 | 0.24 | 5591 |
| Prosocial SDQ | 0.11 | 0.52 | 0.05 | 0.02 | 0.03 | 0.27 | 0.63 | 0.07 | 0.3 | 5591 |
| **Teacher Report Age 7** |  |  |  |  |  |  |  |  |  |  |
|  | Ac | As | Cc | Cs | Ec | Es | Tot-H2 | Tot-C2 | Tot-E2 | N |
| Hyperactivity SDQ | 0.26 | 0.44 | 0 | 0 | 0.07 | 0.23 | 0.7 | 0 | 0.3 | 4608 |
| Conduct SDQ | 0.37 | 0.31 | 0 | 0 | 0.09 | 0.22 | 0.69 | 0 | 0.31 | 4610 |
| Peer Problems SDQ | 0.21 | 0.43 | 0 | 0 | 0.05 | 0.3 | 0.64 | 0 | 0.36 | 4602 |
| Emotional Problems SDQ | 0.02 | 0.48 | 0 | 0 | 0.01 | 0.49 | 0.5 | 0 | 0.5 | 4591 |
| APSD Psychopathy= Narcissism + CU-Traits | 0.7 | 0.04 | 0 | 0 | 0.18 | 0.08 | 0.74 | 0 | 0.26 | 4591 |
| Asd = Social + Nonsocial | 0.23 | 0.47 | 0 | 0 | 0.06 | 0.24 | 0.7 | 0 | 0.3 | 4601 |
| Prosocial SDQ | 0.53 | 0.21 | 0 | 0 | 0.13 | 0.13 | 0.73 | 0 | 0.27 | 4603 |
| **Child Report Age 9** |  |  |  |  |  |  |  |  |  |  |
|  | Ac | As | Cc | Cs | Ec | Es | Tot-H2 | Tot-C2 | Tot-E2 | N |
| Hyperactivity SDQ | 0.17 | 0.17 | 0.06 | 0 | 0.08 | 0.52 | 0.34 | 0.06 | 0.6 | 2635 |
| Conduct SDQ | 0.21 | 0.17 | 0.08 | 0 | 0.1 | 0.44 | 0.38 | 0.08 | 0.54 | 2631 |
| Peer Problems SDQ | 0.17 | 0.17 | 0.06 | 0 | 0.08 | 0.51 | 0.34 | 0.06 | 0.6 | 2621 |
| Emotional Problems SDQ | 0.19 | 0.19 | 0.07 | 0 | 0.09 | 0.46 | 0.38 | 0.07 | 0.56 | 2633 |
| Cast= Social + Nonsocial | 0.22 | 0.16 | 0.08 | 0 | 0.11 | 0.43 | 0.38 | 0.08 | 0.54 | 2556 |
| Prosocial SDQ | 0.04 | 0.33 | 0.01 | 0.02 | 0.02 | 0.58 | 0.37 | 0.04 | 0.59 | 2637 |
| **Parent Report Age 9** |  |  |  |  |  |  |  |  |  |  |
|  | Ac | As | Cc | Cs | Ec | Es | Tot-H2 | Tot-C2 | Tot-E2 | N |
| Hyperactivity SDQ | 0.2 | 0.32 | 0.15 | 0 | 0.04 | 0.3 | 0.51 | 0.15 | 0.34 | 2671 |
| Conduct SDQ | 0.31 | 0.23 | 0.23 | 0.05 | 0.07 | 0.11 | 0.54 | 0.28 | 0.18 | 2673 |
| Peer Problems SDQ | 0.1 | 0.52 | 0.07 | 0 | 0.02 | 0.28 | 0.62 | 0.07 | 0.31 | 2673 |
| Emotional Problems SDQ | 0.08 | 0.45 | 0.06 | 0.02 | 0.02 | 0.37 | 0.53 | 0.08 | 0.39 | 2673 |
| Cast | 0.16 | 0.54 | 0.12 | 0 | 0.04 | 0.15 | 0.7 | 0.12 | 0.18 | 2675 |
| APSD Psychopathy | 0.4 | 0.13 | 0.3 | 0 | 0.09 | 0.08 | 0.53 | 0.3 | 0.17 | 2676 |
| Aggression | 0.24 | 0.28 | 0.18 | 0.14 | 0.06 | 0.1 | 0.52 | 0.32 | 0.16 | 2666 |
| Prosocial SDQ | 0.09 | 0.53 | 0.07 | 0.14 | 0.02 | 0.16 | 0.61 | 0.21 | 0.18 | 2675 |
| **Teacher Report Age 9** |  |  |  |  |  |  |  |  |  |  |
|  | Ac | As | Cc | Cs | Ec | Es | Tot-H2 | Tot-C2 | Tot-E2 | N |
| Hyperactivity SDQ | 0.33 | 0.27 | 0.02 | 0 | 0.12 | 0.27 | 0.59 | 0.02 | 0.39 | 2217 |
| Conduct SDQ | 0.47 | 0.14 | 0.03 | 0 | 0.17 | 0.2 | 0.61 | 0.03 | 0.36 | 2221 |
| Peer Problems SDQ | 0.16 | 0.34 | 0.01 | 0.07 | 0.06 | 0.37 | 0.5 | 0.08 | 0.42 | 2229 |
| Emotional Problems SDQ | 0.03 | 0.46 | 0 | 0 | 0.01 | 0.5 | 0.49 | 0 | 0.51 | 2225 |
| Cast | 0.26 | 0.43 | 0.02 | 0 | 0.09 | 0.2 | 0.69 | 0.02 | 0.29 | 2228 |
| APSD Psychopathy | 0.64 | 0.02 | 0.04 | 0 | 0.23 | 0.08 | 0.66 | 0.04 | 0.31 | 2229 |
| Aggression | 0.45 | 0.2 | 0.03 | 0 | 0.16 | 0.17 | 0.65 | 0.03 | 0.33 | 2216 |
| Prosocial SDQ | 0.29 | 0.22 | 0.02 | 0.1 | 0.1 | 0.26 | 0.51 | 0.12 | 0.37 | 2226 |
| **Parent Report Age 12** |  |  |  |  |  |  |  |  |  |  |
|  | Ac | As | Cc | Cs | Ec | Es | Tot-H2 | Tot-C2 | Tot-E2 | N |
| Prosocial | 0.1 | 0.6 | 0.05 | 0.02 | 0.02 | 0.2 | 0.7 | 0.07 | 0.22 | 4654 |
| Conduct | 0.31 | 0.27 | 0.17 | 0.04 | 0.07 | 0.14 | 0.58 | 0.21 | 0.21 | 4645 |
| Depression | 0.16 | 0.33 | 0.09 | 0.04 | 0.04 | 0.34 | 0.49 | 0.13 | 0.38 | 4644 |
| Peer Problems | 0.15 | 0.48 | 0.08 | 0 | 0.03 | 0.27 | 0.62 | 0.08 | 0.3 | 4644 |
| Emotional Problems | 0.11 | 0.38 | 0.06 | 0 | 0.02 | 0.42 | 0.49 | 0.06 | 0.45 | 4644 |
| Cast | 0.2 | 0.44 | 0.11 | 0 | 0.04 | 0.2 | 0.65 | 0.11 | 0.24 | 4743 |
| Conners | 0.31 | 0.34 | 0.17 | 0 | 0.07 | 0.11 | 0.66 | 0.17 | 0.17 | 4652 |
| APSD | 0.37 | 0.26 | 0.2 | 0 | 0.08 | 0.09 | 0.63 | 0.2 | 0.17 | 4643 |
| **Teacher Report Age 12** |  |  |  |  |  |  |  |  |  |  |
|  | Ac | As | Cc | Cs | Ec | Es | Tot-H2 | Tot-C2 | Tot-E2 | N |
| Prosocial | 0.35 | 0.22 | 0 | 0.02 | 0.13 | 0.28 | 0.57 | 0.02 | 0.41 | 3812 |
| Conduct | 0.46 | 0.13 | 0 | 0 | 0.18 | 0.23 | 0.59 | 0 | 0.41 | 3842 |
| Peer Problems | 0.12 | 0.44 | 0 | 0 | 0.04 | 0.4 | 0.56 | 0 | 0.44 | 3842 |
| Emotional Problems | 0.04 | 0.39 | 0 | 0 | 0.02 | 0.55 | 0.44 | 0 | 0.56 | 3836 |
| CAST | 0.22 | 0.35 | 0 | 0 | 0.08 | 0.34 | 0.57 | 0 | 0.43 | 3811 |
| APSD | 0.61 | 0.06 | 0 | 0 | 0.23 | 0.1 | 0.67 | 0 | 0.33 | 3828 |
| **Child Report Age 12** |  |  |  |  |  |  |  |  |  |  |
|  | Ac | As | Cc | Cs | Ec | Es | Tot-H2 | Tot-C2 | Tot-E2 | N |
| Prosocial | 0.01 | 0.41 | 0 | 0 | 0.01 | 0.57 | 0.42 | 0 | 0.58 | 4631 |
| Conduct | 0.14 | 0.24 | 0.04 | 0 | 0.1 | 0.48 | 0.37 | 0.04 | 0.59 | 4631 |
| Peer Problems | 0.16 | 0.2 | 0.05 | 0 | 0.12 | 0.47 | 0.36 | 0.05 | 0.59 | 4631 |
| Emotional Problems | 0.23 | 0.1 | 0.07 | 0 | 0.17 | 0.42 | 0.33 | 0.07 | 0.6 | 4631 |
| Depression | 0.3 | 0.01 | 0.1 | 0.02 | 0.23 | 0.33 | 0.32 | 0.12 | 0.56 | 4649 |
| **Parent Report Age 16** |  |  |  |  |  |  |  |  |  |  |
|  | Ac | As | Cc | Cs | Ec | Es | Tot-H2 | Tot-C2 | Tot-E2 | N |
| Prosocial | 0.22 | 0.33 | 0.1 | 0.18 | 0.05 | 0.1 | 0.56 | 0.28 | 0.15 | 4011 |
| Conduct | 0.28 | 0.31 | 0.13 | 0 | 0.6 | 0.2 | 0.6 | 0.13 | 0.26 | 4011 |
| Depression | 0.15 | 0.3 | 0.07 | 0 | 0.03 | 0.43 | 0.46 | 0.7 | 0.12 | 4011 |
| Cast | 0.14 | 0.66 | 0.06 | 0 | 0.02 | 0.09 | 0.81 | 0.6 | 0.12 | 4010 |
| Cu | 0.31 | 0.27 | 0.14 | 0.07 | 0.6 | 0.12 | 0.58 | 0.22 | 0.18 | 4010 |
| Conners | 0.23 | 0.44 | 0.1 | 0 | 0.04 | 0.16 | 0.68 | 0.1 | 0.21 | 4010 |
| Arbq | 0.15 | 0.34 | 0.07 | 0.07 | 0.03 | 0.32 | 0.49 | 0.14 | 0.35 | 3995 |
| **Child Report Age 16** |  |  |  |  |  |  |  |  |  |  |
|  | Ac | As | Cc | Cs | Ec | Es | Tot-H2 | Tot-C2 | Tot-E2 | N |
| Prosocial | 0.01 | 0.37 | 0.00 | 0.00 | 0.01 | 0.62 | 0.38 | 0.00 | 0.62 | 3999 |
| Conduct | 0.08 | 0.26 | 0.00 | 0.00 | 0.06 | 0.61 | 0.33 | 0.00 | 0.67 | 3998 |
| Peer Problems | 0.17 | 0.25 | 0.00 | 0.00 | 0.12 | 0.46 | 0.41 | 0.00 | 0.59 | 3999 |
| Emotional Problems | 0.35 | 0.06 | 0.00 | 0.00 | 0.26 | 0.33 | 0.41 | 0.00 | 0.59 | 3999 |
| Depression | 0.34 | 0.06 | 0.00 | 0.00 | 0.25 | 0.35 | 0.40 | 0.00 | 0.60 | 4000 |
| Cast | 0.13 | 0.35 | 0.00 | 0.00 | 0.10 | 0.42 | 0.48 | 0.00 | 0.52 | 4002 |

Table S2: Model fit statistics for common pathway twin models of childhood psychopathology. Note that Sat= saturated model; CPACE = the common pathway ACE model

| **Model** | **base** | **comparison** | **ep** | **minus2LL** | **df** | **AIC** | **diffLL** | **diffdf** | **p** |
| --- | --- | --- | --- | --- | --- | --- | --- | --- | --- |
| Age 7 parent report | Sat | CPACE | 38 | 198398.6 | 77905 | 42588.63 | 8872.522 | 201 | 0.00 |
| Age 7 teacher report | Sat | CPACE | 38 | 153092 | 64219 | 24654 | 8562.541 | 201 | 0.00 |
| Age 9 self report | Sat | CPACE | 33 | 83002.4 | 31375 | 20252.4 | 1032.731 | 148 | 0.00 |
| Age 9 parent report | Sat | CPACE | 43 | 82157.2 | 35470 | 11217.2 | 4209.129 | 262 | 0.00 |
| Age 9 teacher report | Sat | CPACE | 43 | 101865.31 | 42701 | 16463.31 | 3960.66 | 262 | 0.00 |
| Age12 self report | Sat | CPACE | 28 | 121208.10 | 46270.00 | 28668.05 | 966.53 | 103.00 | 0.00 |
| Age 12 parent report | Sat | CPACE | 43 | 178367.50 | 74408.00 | 29551.50 | 5872.72 | 262.00 | 0.00 |
| Age 12 teacher report | Sat | CPACE | 33 | 113280.20 | 45806.00 | 21668.24 | 4135.12 | 148.00 | 0.00 |
| Age16 self report | Sat | CPACE | 48 | 170085.10 | 65525.00 | 39035.12 | 4212.57 | 331.00 | 0.00 |
| Age 16 parent report | Sat | CPACE | 38 | 134663.40 | 55982.00 | 22699.37 | 6624.52 | 201.00 | 0.00 |

Supplementary Table S3: Loadings on first principal components of psychopathology measures for each age and rater.

| Parent report age 7 | |
| --- | --- |
| Hyperactivity SDQ | 0.34 |
| Conduct SDQ | 0.42 |
| Peer problems SDQ | 0.38 |
| Emotional problems SDQ | 0.29 |
| APSD psychopathy = Narcissism + CU-traits | 0.41 |
| ASD = Social + Nonsocial | 0.45 |
| Prosocial SDQ | 0.32 |
| Teacher report age 7 | |
| Hyperactivity SDQ | 0.35 |
| Conduct SDQ | 0.40 |
| Peer problems SDQ | 0.39 |
| Emotional problems SDQ | 0.22 |
| APSD psychopathy= Narcissism + CU-traits | 0.45 |
| ASD = Social + Nonsocial | 0.40 |
| Prosocial SDQ | 0.40 |
| Child report age 9 | |
| Hyperactivity SDQ | 0.42 |
| Conduct SDQ | 0.45 |
| Peer problems SDQ | 0.42 |
| Emotional problems SDQ | 0.43 |
| CAST= Social + Nonsocial | 0.46 |
| Prosocial SDQ | 0.21 |
| Parent report age 9 | |
| Hyperactivity SDQ | 0.36 |
| Conduct SDQ | 0.41 |
| Peer problems SDQ | 0.32 |
| Emotional problems SDQ | 0.30 |
| CAST | 0.38 |
| APSD psychopathy | 0.44 |
| Aggression | 0.34 |
| Prosocial SDQ | 0.22 |
| Teacher report age 9 | |
| Hyperactivity SDQ | 0.36 |
| Conduct SDQ | 0.40 |
| Peer problems SDQ | 0.32 |
| Emotional problems SDQ | 0.17 |
| CAST | 0.38 |
| APSD psychopathy | 0.46 |
| Aggression | 0.31 |
| Prosocial SDQ | 0.35 |

| Child report age 12 |  |
| --- | --- |
| Prosocial | 0.16 |
| Peer Problems SDQ | 0.47 |
| Emotional Problems SDQ | 0.5 |
| Conduct | 0.48 |
| Depression | 0.54 |
| Parent report age 12 |  |
| Conners | 0.5 |
| Conduct SDQ | 0.5 |
| CU-traits | 0.32 |
| Emotional problems SDQ | 0.43 |
| Depression | 0.47 |
| Teacher report age 12 |  |
| Conduct SDQ | 0.44 |
| Peer problems SDQ | 0.38 |
| Emotional problems SDQ | 0.26 |
| CAST | 0.44 |
| APSD psychopathy | 0.47 |
| Prosocial SDQ | 0.43 |
| Child report age 16 |  |
| Prosocial | 0.07 |
| Peer Problems SDQ | 0.32 |
| Emotional Problems SDQ | 0.49 |
| Conduct | 0.4 |
| Depression | 0.47 |
| CAST | 0.39 |
| CASI | 0.41 |
| Parent report age 16 |  |
| Conduct SDQ | 0.4 |
| Conners | 0.38 |
| CAST | 0.36 |
| ARBQ psychopathy | 0.36 |
| Prosocial SDQ | 0.36 |
| Depression | 0.34 |
| CU | 0.4 |

*Supplementary Table S4.* Variance explained by first principal components (phenotypic p factors) for each age and rater, plus the sample size for each 1^st^ PC. P/T/C= parent/teacher/child ratings; 7/9/12/16 = age.

| **Age/rater** | **R^2^ by 1st PC** | **N** |
| --- | --- | --- |
| P7 | 0.40 | 4109 |
| T7 | 0.50 | 3435 |
| C9 | 0.42 | 2074 |
| P9 | 0.45 | 2157 |
| T9 | 0.48 | 1594 |
| C12 | 0.46 | 4490 |
| P12 | 0.45 | 3227 |
| T12 | 0.50 | 2146 |
| C16 | 0.42 | 1391 |
| P16 | 0.46 | 3258 |

*Supplementary Table S5*. Association statistics for polygenic p across phenotypic p measures.

| **Outcome** | **beta** | **se** | **p-value** | **r squared** | **df residual** |
| --- | --- | --- | --- | --- | --- |
| P age 7 | 0.083554 | 0.015365 | **5.70E-08** | 0.007149 | 4107 |
| P age 9 | 0.082633 | 0.021656 | **0.00014** | 0.006711 | 2155 |
| P age 12 | 0.085961 | 0.017255 | **6.63E-07** | 0.007637 | 3225 |
| P age 16 | 0.078936 | 0.017206 | **4.65E-06** | 0.006423 | 3256 |
| T age 7 | 0.097842 | 0.016814 | **6.47E-09** | 0.009767 | 3433 |
| T age 9 | 0.089132 | 0.025625 | **0.000518** | 0.007542 | 1592 |
| T age 12 | 0.055183 | 0.020751 | 0.007889 | 0.003288 | 2144 |
| C age 9 | 0.066584 | 0.022225 | **0.002769** | 0.004313 | 2072 |
| C age 12 | 0.065225 | 0.014811 | **1.09E-05** | 0.004303 | 4488 |
| C age 16 | 0.068554 | 0.026306 | 0.009258 | 0.005750 | 1389 |

**Bold**. Significant after Bonferroni correction (alpha 0.05/10).

**Note.** P = parent rated , T = teacher rated, C = child-rated.
